# GWAS for serum galactose-deficient IgA1 implicates critical genes of the *O*-glycosylation pathway

**DOI:** 10.1101/076414

**Authors:** Krzysztof Kiryluk, Yifu Li, Zina Moldoveanu, Hitoshi Suzuki, Colin Reily, Ping Hou, Jingyuan Xie, Nikol Mladkova, Sindhuri Prakash, Clara Fischman, Samantha Shapiro, Robert A. LeDesma, Drew Bradbury, Iuliana Ionita-Laza, Frank Eitner, Thomas Rauen, Nicolas Maillard, Francois Berthoux, Jürgen Floege, Nan Chen, Hong Zhang, Francesco Scolari, Robert J. Wyatt, Bruce A. Julian, Ali G. Gharavi, Jan Novak

**Author notes:** Corresponding Author: Krzysztof Kiryluk M.D., M.S. Department of Medicine Division of Nephrology Columbia University 1150 St Nicholas Ave Russ Berrie Pavilion #412 New York, NY 10032.

## Abstract

Aberrant *O*-glycosylation of serum immunoglobulin A1 (IgA1) represents a heritable pathogenic defect in IgA nephropathy, the most common form of glomerulonephritis worldwide, but specific genetic factors involved in its determination are not known. We performed a quantitative GWAS for serum levels of galactose-deficient IgA1 (Gd-IgA1) in 2,633 subjects of European and East Asian ancestry and discovered two genome-wide significant loci, in *C1GALT1* (rs13226913, *P* = 3.2 × 10^−11^) and *C1GALT1C1* (rs5910940, *P* = 2.7 × 10^−8^). These genes encode molecular partners essential for enzymatic *O*-glycosylation of IgA1. We demonstrated that these two loci explain approximately 7% of variability in circulating Gd-IgA1 in Europeans, but only 2% in East Asians. Notably, the Gd-IgA1-increasing allele of rs13226913 is common in Europeans, but rare in East Asians. Moreover, rs13226913 represents a strong cis-eQTL for *C1GALT1,* which encodes the key enzyme responsible for the transfer of galactose to *O*-linked glycans on IgA1. By *in vitro* siRNA knock-down studies, we confirmed that mRNA levels of both *C1GALT1* and *C1GALT1C1* determine the rate of secretion of Gd-IgA1 in IgA1-producing cells. Our findings provide novel insights into the genetic regulation of *O*-glycosylation and are relevant not only to IgA nephropathy, but also to other complex traits associated with *O*-glycosylation defects, including inflammatory bowel disease, hematologic disease, and cancer.

**Author Summary:** *O*-glycosylation is a common type of post-translational modification of proteins; specific abnormalities in the mechanism of *O*-glycosylation have been implicated in cancer, inflammatory and blood diseases. However, the molecular basis of abnormal *O*-glycosylation in these complex disorders is not known. We studied the genetic basis of defective *O*-glycosylation of serum Immunoglobulin A1 (IgA1), which represents the key pathogenic defect in IgA nephropathy, the most common form of primary glomerulonephritis worldwide. We report our results of the first genome-wide association study for this trait using serum assays in 2,633 individuals of European and East Asian ancestry. In our genome scan, we observed two significant signals with large effects, on chromosomes 7p21.3 and Xq24, jointly explaining about 7% of trait variability. These signals implicate two genes that encode molecular partners essential for enzymatic *O*-glycosylation of IgA1 and mucins, and represent potential new targets for therapy.

## Introduction

*N*- and *O*-glycosylation is a fundamental post-translational modification of proteins in mammalian cells. Abnormalities in glycosylation have been linked to a broad range of human diseases, including neurologic disorders, immune-mediated and inflammatory diseases as well as cancer. Protein glycosylation is mediated by a large family of enzymes that have cell and tissue specific activity, and can generate highly diverse glycan structures that are important for signaling, cell-cell and cell-matrix interactions. The combinatorial possibilities of glycan structures imparted by the large number of glycosylation enzymes complicate a systematic analysis of protein glycosylation patterns and identification of critical steps involved in the activity, concentration, and regulation in any given cell or tissue. In such setting, genetic studies of congenital defects of glycosylation in humans have provided significant insight into non-redundant regulatory nodes in this pathway. The majority of these Mendelian disorders arise from loss of function mutations that severely perturb protein glycosylation across a range of tissues and produce a wide range of organ dysfunction in early life. However, less pronounced abnormalities in protein glycosylation have also been detected in complex disorders such as autoimmunity and cancer, suggesting that more subtle defects in this pathway can have important consequences for human health.

IgA nephropathy (IgAN), the most common cause of glomerulonephritis and a common cause of kidney failure worldwide, is a prototypical example of an immune-mediated disorder characterized by abnormal glycosylation^1^. In humans, the hinge region heavy chains of immunoglobulin A1 (IgA1) are *O*-glycosylated with a variety of glycoforms in circulation. In healthy individuals, the prevailing glycoforms include the *N*-acetylgalactosamine (GalNAc)-galactose disaccharide and its sialylated forms. In IgAN, galactose-deficient IgA1 (Gd-IgA1) glycoforms are significantly more abundant compared to healthy controls^2^. These under-galactosylated glycoforms are secreted by IgA1-producing cells while galactosylation of other circulating *O*-glycosylated proteins is preserved, suggesting a specific defect within IgA1-producing cells^3^. The pathogenetic mechanism of IgAN involves an autoimmune response resulting in production of IgA or IgG autoantibodies against circulating Gd-IgA1, and formation of immune complexes (Gd-IgA1 complexed with IgA/IgG autoantibody) that deposit in the kidney and cause tissue injury^1,4^. Consistent with this mechanism, Gd-IgA1 is the predominant glycoform in circulating immune complexes and in the glomerular immune-deposits in patients with IgAN^5-8^ and elevated serum levels of Gd-IgA1 (auto-antigen) and anti-glycan antibodies (auto-antibody) are associated with more aggressive disease and accelerated progression to end-stage kidney failure^9,10^.

The design of a simple lectin-based ELISA assay, using a GalNAc-specific lectin from *Helix aspersa* (HAA), enables high-throughput screening of sera to quantify the levels of circulating Gd-IgA1^2^. Using this assay, we have demonstrated that the serum levels of Gd-IgA1 represent a normally distributed quantitative trait in healthy populations, but up to two thirds of IgAN patients have levels above the 95^th^ percentile for healthy controls. Examining family members of probands with familial and sporadic forms of IgAN, we also showed that elevated serum Gd-IgA1 levels segregate independently of total IgA levels and have high heritability (estimated at 50-70%) ^11,12^. Moreover, many healthy family members exhibited very high Gd-IgA1 levels, identifying elevated Gd-IgA1 as a heritable risk factor that precedes the development of IgAN.

To date, multiethnic genome-wide association studies involving over 20,000 individuals have identified 15 risk loci predisposing to IgAN, highlighting the importance of innate and adaptive immunity in this disorder. Power analyses indicated that discovery of additional risk loci using the case control design will require significant expansion in sample size. However, a systematic analysis of quantitative endophenotypes that are linked to disease pathogenesis, such as Gd-IgA1, has not been conducted to date and may provides the opportunity to discover additional pathogenic pathways using a smaller sample size. In this study, we performed the first GWAS for serum Gd-IgA1 levels, and successfully mapped new loci with surprisingly large contributions to the heritability of the circulating level of Gd-IgA1 independently of IgA levels.

## Results

In order to test if serum levels of Gd-IgA1 remain stable over time, we first performed measurements of total serum immunoglobulin levels along with Gd-IgA1 levels at baseline and at four years of follow-up in 32 individuals of European ancestry followed longitudinally (Figure 1). While total IgG and IgA levels varied with time, Gd-IgA1 levels (normalized for total IgA) remained remarkably stable over a 4-year period (r^2^ = 0.92, P = 1.8 × 10^−13^), demonstrating that *O*-glycosylation of IgA1 is minimally affected by random environmental factors.

**Figure 1:**
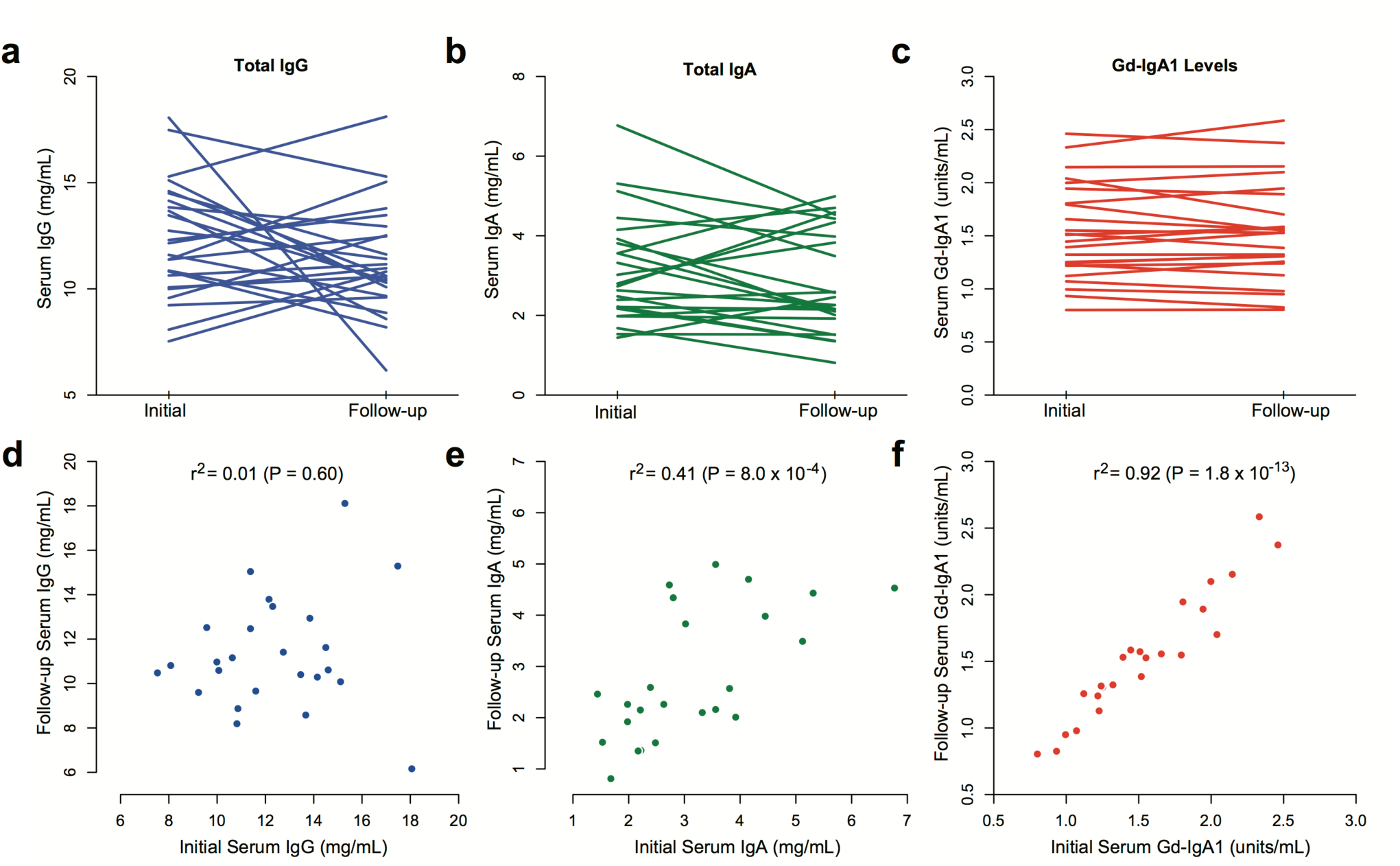
Longitudinal measurements of serum immunoglobulin levels and Gd-IgA1 levels over 4 years of follow-up. Initial and follow-up levels of **(a)** total serum IgG, **(b)** total serum IgA, and **(c)** serum Gd-IgA1 normalized to total serum IgA. Panels **(d, e, f)** represent scatter plots of values at initial visit (x-axis) and 4-year follow-up visit (y-axis). P-values correspond to the Pearson’s test of correlation; r2: squared correlation coefficient.

We next used HAA lectin-based ELISA to analyze single time-point sera of 1,195 individuals in our discovery cohorts composed of 950 individuals of East Asian ancestry (483 biopsy-documented IgAN cases and 467 controls) and 245 individuals of European ancestry (141 biopsy-documented IgAN cases and 104 controls, Table 1). As previously reported, serum Gd-IgA1 levels were positively correlated with age (East Asians r = 0.13, P = 8.9×10^−5^; Europeans r = 0.15, P = 1.7×10^−2^) and total IgA levels (East Asians r = 0.75, P < 2.2×10^−16^; Europeans r = 0.56, P < 2.2×10^−16^), but were independent of gender (P >0.05). In both cohorts, Gd-IgA1 levels were also significantly higher in IgAN cases compared to controls independently of age and total IgA levels (adjusted P < 2.2×10^−16^ in each individual cohort), providing a large-scale replication of prior findings.

**Table 1.**
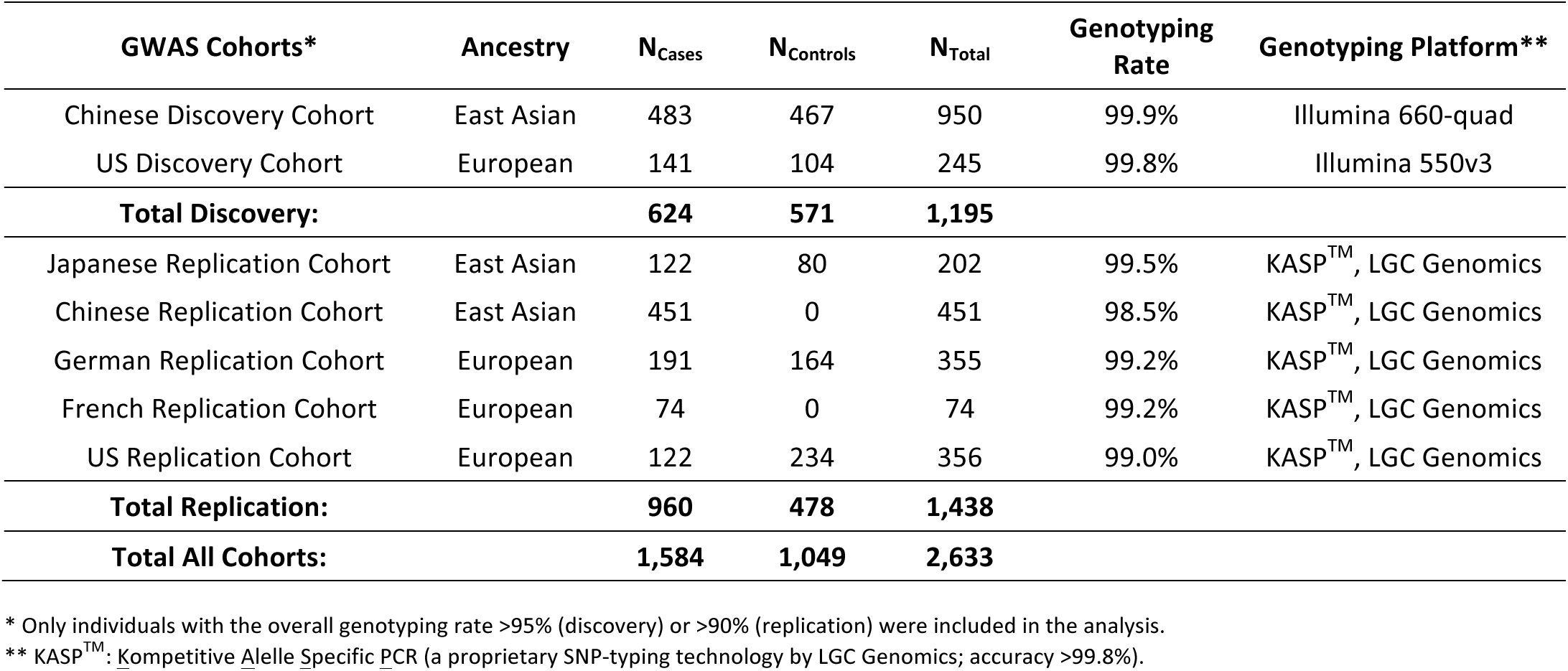
Study Cohorts: the final numbers of cases and controls by cohort after implementation of all quality control filters.

| **GWAS Cohorts\*** | **Ancestry** | **N<sub>Cases</sub>** | **N<sub>Controls</sub>** | **N<sub>Total</sub>** | **Genotyping Rate** | **Genotyping Platform\*\*** |
| --- | --- | --- | --- | --- | --- | --- |
| Chinese Discovery Cohort | East Asian | 483 | 467 | 950 | 99.9% | Illumina 660-quad |
| US Discovery Cohort | European | 141 | 104 | 245 | 99.8% | Illumina 550v3 |
| **Total Discovery:** |  | **624** | **571** | **1,195** |  |  |
| Japanese Replication Cohort | East Asian | 122 | 80 | 202 | 99.5% | KASP<sup>TM</sup>, LGC Genomics |
| Chinese Replication Cohort | East Asian | 451 | 0 | 451 | 98.5% | KASP<sup>TM</sup>, LGC Genomics |
| German Replication Cohort | European | 191 | 164 | 355 | 99.2% | KASP<sup>TM</sup>, LGC Genomics |
| French Replication Cohort | European | 74 | 0 | 74 | 99.2% | KASP<sup>TM</sup>, LGC Genomics |
| US Replication Cohort | European | 122 | 234 | 356 | 99.0% | KASP<sup>TM</sup>, LGC Genomics |
| **Total Replication:** |  | **960** | **478** | **1,438** |  |  |
| **Total All Cohorts:** |  | **1,584** | **1,049** | **2,633** |  |  |

\* Only individuals with the overall genotyping rate >95% (discovery) or >90% (replication) were included in the analysis.
\*\* KASP<sup>TM</sup>: Kompetitive Allele Specific PCR (a proprietary SNP-typing technology by LGC Genomics; accuracy >99.8%).

We next performed a GWAS for serum levels of Gd-IgA1 in these cohorts with and without adjustment for total IgA levels. For genome-wide analysis, we used a linear model with individual SNPs coded as additive genetic predictors, and the outcome defined as standardized residuals of serum Gd-IgA1 after normalization and additional adjustment for case/control status, age, ancestry and cohort membership (see Methods). Each ethnicity-defined discovery cohort was analyzed separately and the results were meta-analyzed to prioritize top signals for follow-up. With this approach, we observed minimal genomic inflation in the combined genome-wide analyses (*λ*=1.01), indicating negligible effect of population stratification.

We first examined potential associations with known IgAN susceptibility loci, but found no statistically significant or suggestive signals between Gd-IgA1 levels and known IgAN risk alleles (**Supplemental Table 1**). In addition, we found no association between the global polygenetic risk score for IgAN, which captures the combined effect of all IgAN risk loci, and Gd-IgA1 levels. We also did not detect any associations of Gd-IgA1 levels with loci previously linked to variation in total IgA levels^13–15^, IgA deficiency^16^ or *N*- glycosylation of IgG^17^. At the same time, we replicated previously reported association of total IgA with *ELL2* (rs56219066, P=8.5×10^−3^)^14^, confirming that genetic regulation of IgA levels is distinct from Gd-IgA1 levels. These data thus indicated the presence of yet undiscovered loci controlling variation in Gd-IgA1 levels.

We next examined genome-wide distribution of P-values from the discovery stage to identify novel loci associated with Gd-IgA1 levels. Although no signals reached genome-wide significance in the discovery stage, we observed a number of suggestive (P < 5×10^−4^) loci that we followed up in 1,438 additional individuals of East Asian (N=653) and European (N=785) ancestry (**Supplementary Figure 1**). Subsequently, we analyzed all cohorts (N=2,633) jointly to identify genome-wide significant loci (Table 2, **Supplementary Table 2**). Our power calculations demonstrate that our design provides adequate power to detect variants explaining >1.5% of overall trait variance at a genome-wide significant alpha 5×10^−8^ (**Supplementary Table 3**).

**Table 2.** Combined results for significant and suggestive GWAS signals: serum Gd-IgA1 levels were adjusted for age, case-control status, and serum total IgA levels. Chromosome X analyses were stratified by sex.

|  |  |  |  | Discovery Cohorts<br>N=1,195 |  |  | Replication Cohorts<br>N=1,438 |  |  | All Cohorts<br>N=2,633 |  |  | Hetero**<br>P-value | Genes in<br>Locus |
| --- | --- | --- | --- | --- | --- | --- | --- | --- | --- | --- | --- | --- | --- | --- |
| Chr | Position<br>(kb) | SNP | Allele* | Effect | SE | P-value | Effect | SE | P-value | Effect | SE | P-value |  |  |
| 7 | 7213 | rs13226913 | T | 0.20 | 0.05 | 2.2E-04 | 0.23 | 0.04 | 3.0E-08 | 0.22 | 0.03 | 3.2E-11 | 0.43 (NS) | **C1GALT1** |
| 7 | 7239 | rs1008897 | G | 0.19 | 0.06 | 4.6E-04 | 0.22 | 0.04 | 4.6E-07 | 0.21 | 0.03 | 9.1E-10 | 0.93 (NS) | **C1GALT1** |
| X | 119642 | rs5910940 | T | 0.13 | 0.03 | 2.3E-04 | 0.11 | 0.03 | 3.8E-05 | 0.14 | 0.03 | 2.7E-08 | 0.88 (NS) | **C1GALT1C1** |
| X | 119698 | rs2196262 | A | 0.11 | 0.03 | 1.2E-03 | 0.10 | 0.03 | 3.3E-04 | 0.12 | 0.02 | 1.4E-06 | 0.46 (NS) | **C1GALT1C1** |
| 7 | 43345 | rs978056 | G | 0.10 | 0.03 | 1.2E-03 | 0.07 | 0.03 | 7.5E-03 | 0.08 | 0.02 | 3.3E-05 | 0.15 (NS) | **HECW1** |

\* Gd-IgA1-increasing allele is provided as reference; \*\* P-value for the test of heterogeneity of effects

In the combined analysis, two distinct genomic loci, on chromosomes 7p21.3 and Xq24, reached genome-wide significance (Figure 2a). The strongest association was located within a 200-kb interval on chromosome 7p21.3 (Figure 2b), explaining 4% of trait variance in Europeans and ~1% in Asians (**Supplementary Table 4**). The only gene within this locus is *C1GALT1,* encoding core 1 synthase, glycoprotein-*N*-acetylgalactosamine 3-beta-galactosyltransferase 1. The top signal was represented by rs13226913 (*P*=3.2×10^−11^), an intronic SNP within *C1GALT1.* This locus is further supported by rs1008897 (*P*=9.1×10^−10^) in partial LD with rs13226913 (r^2^=0.33, D’=0.91 in Europeans and r^2^=0.52, D’=0.73 in Asians). After mutual conditioning, both SNPs continue to be associated with the phenotype, suggesting a complex pattern of association at this locus (**Supplementary Table 5**).

**Figure 2:**
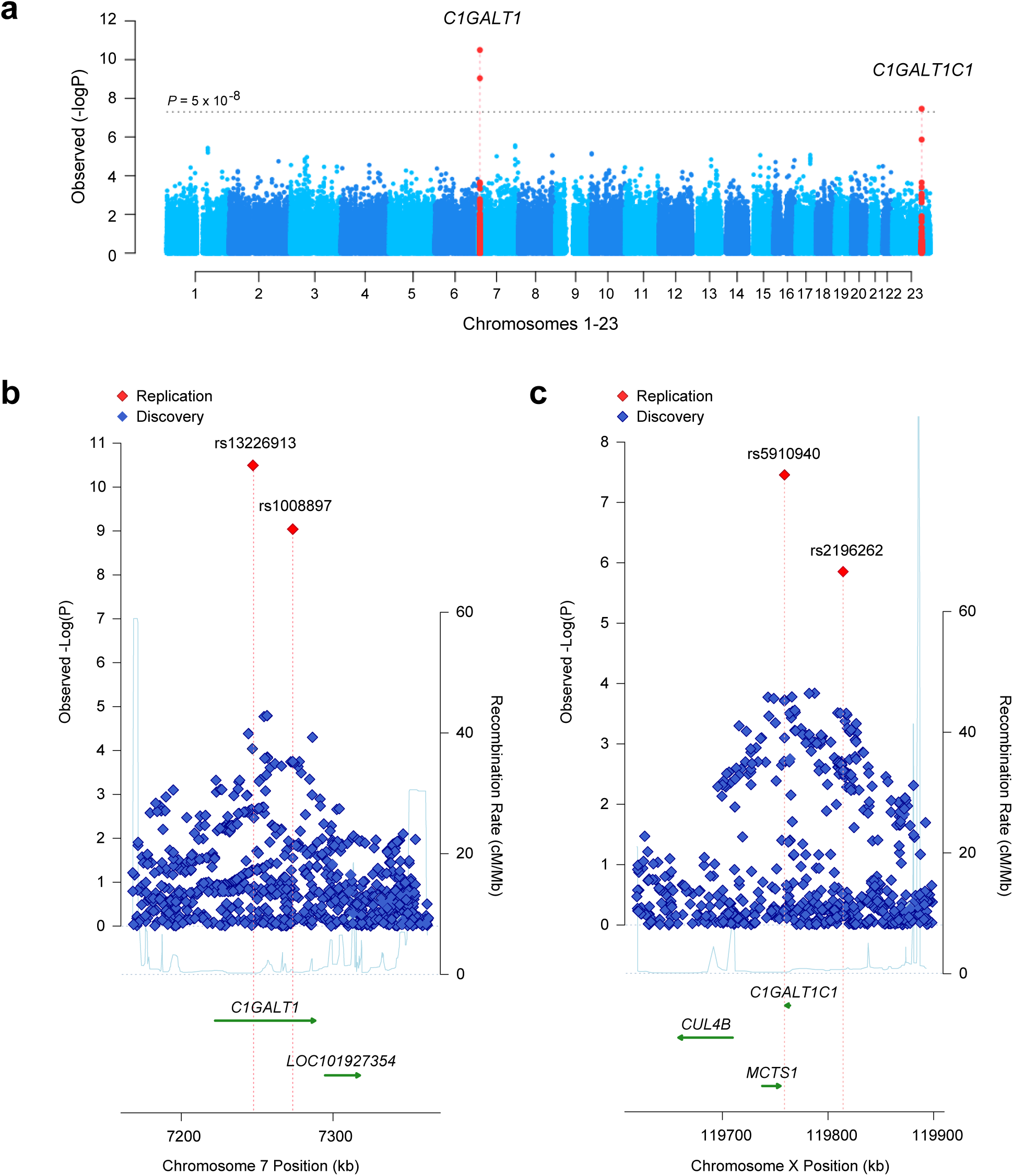
Results of the combined meta-analysis of serum Gd-IgA1 levels in 2,633 individuals of European and Asian ancestry. Manhattan plot **(a)** and regional plots for two distinct genome-wide significant loci, C1GALT1 locus **(b)** and C1GALT1C1 locus **(c)** The x-axis presents physical distance in kilobases (hg18 coordinates), and the y-axis presents -log P values for association statistics. The dotted horizontal line in a represents a genome-wide significance level (P=5×10^−8^). The regional plots contain all genotyped and imputed SNPs in the region meta-analyzed between the discovery and replication cohorts.

The protein encoded by *C1GALT1* generates the common core 1 *O*-glycan structure by transferring galactose (Gal) from UDP-Gal to GalNAc-alpha-1-Ser/Thr. Core 1 is a precursor for *O*-glycans in the hinge region of circulating IgA1, as well as many extended mucin-type *O*-glycans on cell surfaces. In humans, *C1GALT1* is abundantly expressed in IgA1-secreting cells^18^, as well as in EBV-transformed lymphocytes, gastrointestinal tract, lungs, and kidneys^19^. The top SNP, rs13226913, is not in LD with any coding variants, but it perfectly tags several SNPs intersecting the ENCODE and Roadmap enhancers and promoters in immune cells, including EBV-immortalized B cells and primary CD19+ cells (**Supplementary Table 6**). Interrogation of eQTL databases revealed that rs13226913 has a highly significant cis-eQTL effect on *C1GALT1* in peripheral blood (*P*=3.9×10^−23^) with the T allele associated with lower mRNA levels (**Supplementary Table 7**). Consistent with this finding, rs13226913 imparts an additive effect with each T (derived) increasing Gd-IgA1 levels by 0.22 standard deviation units (95%CI: 0.10-0.30).

The second genome-wide significant locus comprises a 100-kb interval on chromosome Xq24 (Figure 2c) and explains an additional 2.7% of the overall trait variance in Europeans and 1.2% in Asians (**Supplementary Table 4**). The top signal at this locus is represented by rs5910940 (*P*=2.7×10^−8^), a SNP 3’ downstream from *C1GALT1C1.* The T (derived) allele increases serum Gd-IgA1 levels by 0.14 standard deviation units per allele (95%CI: 0.11-0.17). Our post-hoc examination of genotypic effects suggests a dominant effect of the rs5910940-T allele in females (dominant model *P*=7.9×10^−9^, **Supplementary Table 8**), although skewed inactivation of chromosome X in IgA1-producing cells could also potentially explain this effect.

*C1GALT1C1* encodes a transmembrane protein that is similar to the core 1 beta1,3-galactosyltransferase 1 encoded by*C1GALT1*. However, its gene product (known as Cosmc) lacks the galactosyltransferase activity, but instead acts as a molecular chaperone required for the folding, stability, and full activity of C1GalT1^20^. *C1GALT1C1* is also ubiquitously expressed in multiple tissues, including IgA1-secreting cells^18^, other blood cells, gastrointestinal tract, kidneys, and lungs^19^. Because sex chromosomes are not included in most eQTL analyses, we were not able to confirm if rs5910940 has an effect on the expression of *C1GALT1C1* based on available datasets. However, rs5910940 tags a 2-bp insertion in the active promoter of *C1GALT1C1* in B-lymphocytes and leukemia cell lines (**Supplementary Table 9**). Considering the known functional dependency of *C1GALT1* and *C1GALT1C1*, we also tested for potential epistasis between these two loci, but did not detect any significant genetic interactions.

Taken together, these data predict an additive regulatory effect of rs13226913 and rs5910940, resulting in lower *C1GALT1* and *C1GALT1C1* expression, and leading to increased production of Gd-IgA1. We next performed siRNA knock-down studies in human cultured IgA1-secreting cell lines to confirm the effect of lower *C1GALT1* and *C1GALT1C1* transcript abundance on the production of Gd-IgA1 (Figure 3). Consistent with the observed genetic effect, *in vitro* knock-down of *C1GALT1* resulted in 30-50% increased production of Gd-IgA1 by the cells derived from IgAN patients (*P*=0.025) as well as healthy controls (*P*=0.011). Similar to *C1GALT1*, *in vitro* siRNA knock-down of *C1GALT1C1* in IgA1-producing cell lines significantly increased the production of Gd-IgA1 in healthy individuals (*P*=0.032) and a similar trend was observed in IgAN patients (*P*=0.066, Figure 3). Consistent with the genetic data, there were no multiplicative effects on Gd-IgA1 production with combined siRNA knock-down in IgA1-secreting lines.

**Figure 3:**
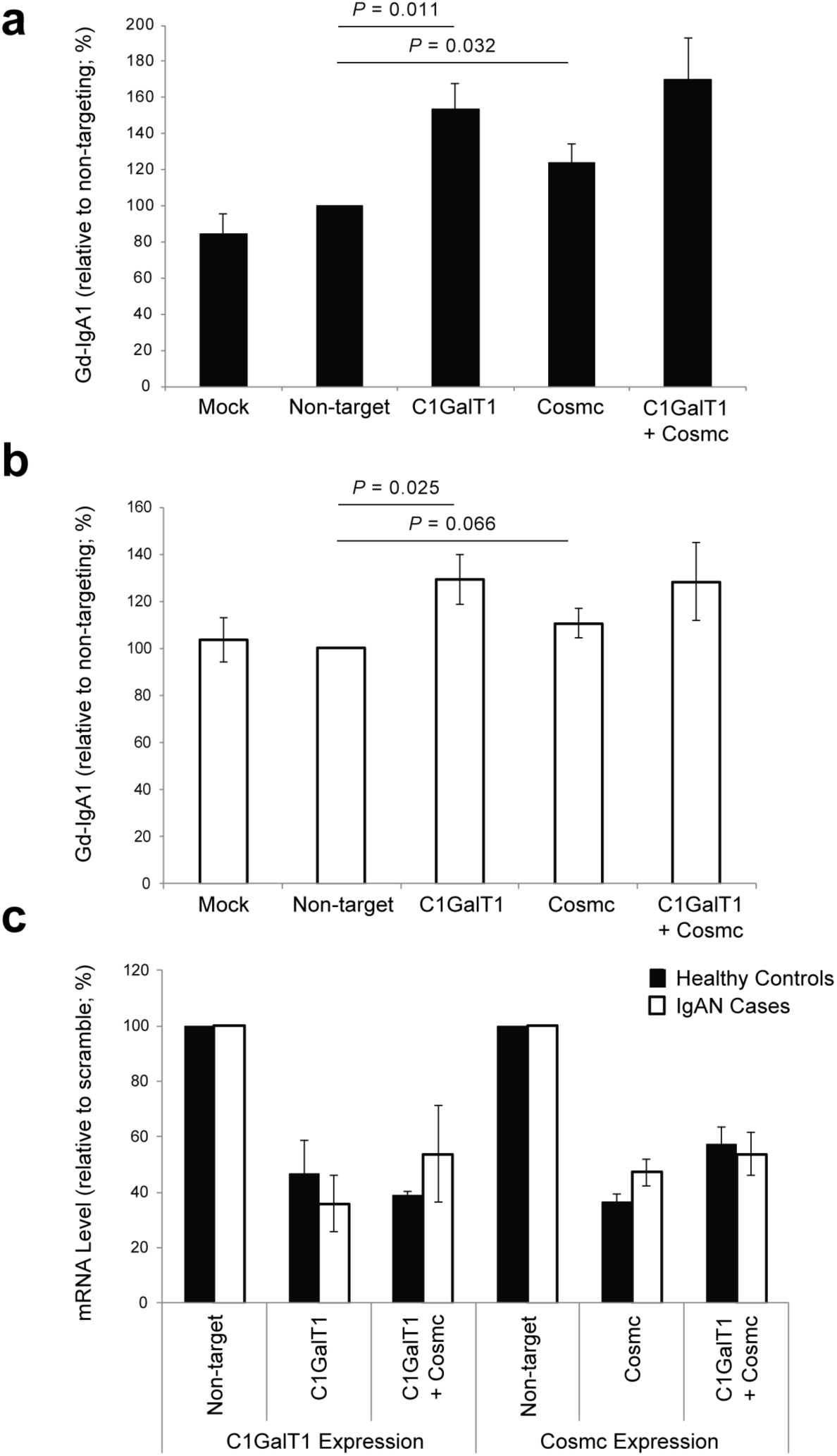
siRNA knock-down of C1GalT1, Cosmc and Cosmc+C1GalT1 in IgA1-secreting cell lines increases Gd-IgA1 production. **(a)** Knock-down in IgA1-secreting cell lines from healthy controls; mock-control (n=5), non-targeting siRNA (n=7), C1GalT1 siRNA (n=5), Cosmc siRNA (n=7), and Cosmc+C1GalT1 siRNA (n=2); **(b)** Knock-down in IgA1-secreting cell lines from IgAN patients; mockcontrol (n=5), non-targeting siRNA (n=7), C1GalT1 (n=5), Cosmc siRNA (n=7), and Cosmc+C1GalT1 siRNA (n=2); **(c)** Relative change in mRNA in IgA1-secreting cell lines after siRNA knock-down of C1GalT1 (n=5), Cosmc (n=7), and Cosmc+C1GalT1 (n=2) compared to non-targeting siRNA control.

Jointly, the newly discovered *C1GALT1* and *C1GALTC1* loci explain up to **7%** of variance in Gd-IgA1 levels in Europeans and **2%** in Asians (**Supplementary Table 4**). Further examination of effect estimates by ethnicity confirms that the European cohorts predominantly drive these associations (**Supplementary Table 10**). Notably, the derived (T) allele of rs13226913 at *C1GALT1* locus is considerably more frequent in Europeans (freq. 47%) compared to Asians (freq. 10%), additionally contributing to the difference in variance explained between ethnicities. Subsequent examination of allelic frequencies in the Human Genome Diversity Panel (Figure 4) confirms that the derived allele of rs13226913 is rare or absent in some Asian populations, while being the predominant (major) allele in Europeans (freq. >50%). In contrast, the T (derived) allele of rs5910940 at *C1GALT1C1* locus is equally frequent in Asian and European populations (freq. ~50%), but nearly fixed in selected African populations. These findings suggest potential involvement of geographically confined selective pressures acting on the loci controlling the *O*-glycosylation process.

**Figure 4:**
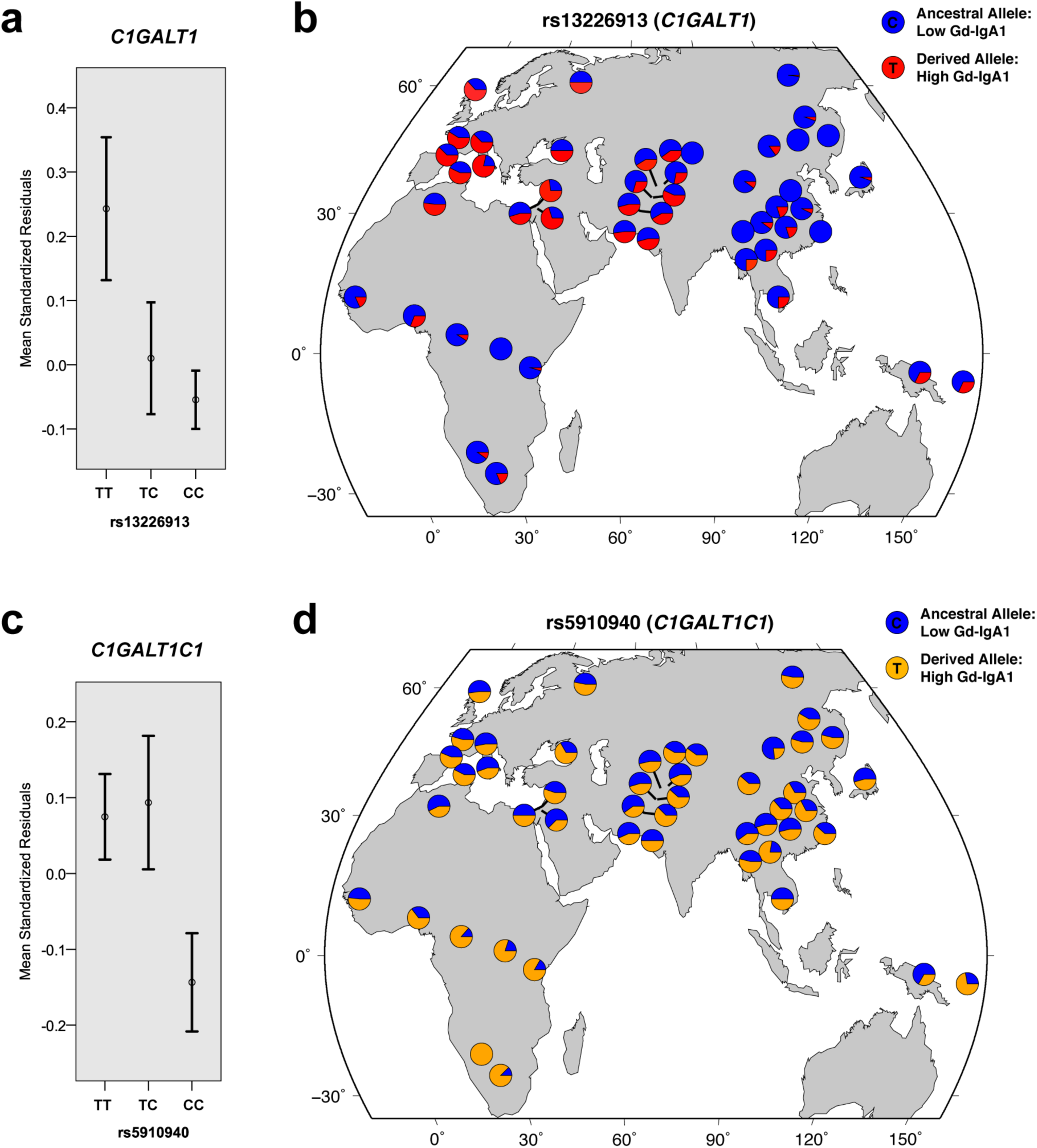
Genotypic effects and worldwide allelic frequency distribution for the two top genome-wide significant loci. (**a**) Mean trait values (+/- standard errors) by rs13226913 genotype at the C1GALT1 locus. (**b**) The distribution of rs13226913 alleles across HGDP populations. (**c**) Mean trait values (+/- standard errors) by rs5910940 genotype at the C1GALT1C1 locus. (**d**) The distribution of rs5910940 alleles across HGDP populations. The allelic distribution plots were modified from the HGDP Selection Browser. The trait values were expressed as standard normal residuals of log-transformed serum Gd-IgA1 levels after adjustment for age, serum total IgA levels, case-control status and cohort membership.

Lastly, we detected additional suggestive signals, including a locus on chromosome 7p13 that warrants further follow-up in larger cohorts (**Supplementary Figure 2a and 2b**). This locus is represented by rs978056 (*P*=3.3×10^−5^), an intronic SNP in *HECW1* (encoding E3 ubiquitin ligase) previously studied in the context of colon and breast cancer (**Supplementary Figure 3**). Based on the analysis of known protein-protein interactions, HECW1 is a second-degree neighbor of C1GalT1 and Cosmc, with ubiquitin C as a common interacting protein (**Supplementary Figure 2c**).

## Discussion

Genetic studies of immune endophenotypes have provided novel insights into the genetic architecture of complex traits and enhanced sub-classification of several autoimmune and inflammatory disorders. The power of immune endophenotypes is best exemplified by recent genetic studies of ANCA titers in vasculitis^21^, IgE levels in asthma^22,23^, or studies of IgG *N*-glycosylation and autoimmunity^17^. Taking a similar approach, we performed the first GWAS for aberrant *O*-glycosylation of IgA1.

Abnormalities in the *O*-glycan synthesis have been linked to several human diseases, including IgA nephropathy, inflammatory bowel disease (IBD), hematologic diseases, and cancer. Mucin-type *O*-glycans produced by epithelial cells are critical for the formation of protective viscous barrier with anti-microbial properties at the mucosal surfaces of the gastrointestinal, urogenital and respiratory systems. Recent studies indicate that proper *O*-glycosylation of mucins is required for intestinal integrity in mice^24,25^ and may play a role in human susceptibility to IBD^26,27^. In addition, *O*-glycosylation can affect the structure and immunogenicity of the modified proteins. For example, defective *O*-glycosylation represents the key pathogenic feature of Tn-syndrome^28^, where acquired enzymatic defect in the addition of galactose to *O*-glycans leads to exposed terminal N-acetylgalactosamine residue (Tn-antigen) on the surface of red blood cells, triggering polyagglutination by naturally occurring anti-Tn antibodies^28^. Moreover, Tn and sialyl-Tn represent oncofetal antigens that are over-expressed in human cancers and may directly influence cancer growth, metastasis and survival, but the exact molecular perturbations that lead to *O*-glycosylation defects in tumor cells are presently not known^29^.

Similar to Tn-syndrome, the pathogenesis of IgA nephropathy involves autoimmune response to Tn antigens. In this case, the Tn-antigen is exposed at the hinge region of IgA1 molecules as a result of aberrant *O*-glycosylation of IgA1 in the endoplasmic reticulum of IgA1-producing cells ^8^. In patients with IgA nephropathy, the galactose-deficient IgA1 (Gd-IgA1) is recognized by circulating anti-Tn antibodies ^4^, leading to the formation of nephritogenic immune complexes ^5–8^. Several independent studies, including in healthy twins and in families with IgA nephropathy, have demonstrated that serum levels of Gd-IgA1 have high heritability, providing high level of support for a genetic determination of this trait and a strong rationale for this study ^11,12,30^.

In this study, we quantified the levels of Gd-IgA1 in sera of 2,633 subjects of European and East Asian ancestry using a simple lectin-based ELISA assay. Using GWAS approach, we discovered two genome-wide significant loci, on chromosomes 7p21.3 and Xq24, both with large effects on circulating levels of Gd-IgA1. The 7p21.3 locus contains *C1GALT1* gene, which encodes human core 1 β1–3-galactosyltransferase (C1GalT1), the key enzyme responsible for the addition of galactose to the Tn antigen. Mice deficient in C1galt1 protein develop thrombocytopenia and kidney disease attributed to defective *O*-glycosylation of cell-surface proteins^31^. Moreover, C1galt1 deficiency in mice results in a defective mucus layer leading to spontaneous colonic inflammation that is dependent on the exposure to intestinal microbiota^24,25^. C1GalT1 requires a molecular chaperone, Cosmc, which ensures that the enzyme is properly folded within the ER; loss of Cosmc activity results in C1GalT1 being degraded in the proteosome^28^. Interestingly, Cosmc is encoded by *C1GALT1C1* residing within our second genome-wide significant locus on chromosome Xq24. We also localized a suggestive locus on chromosome 7p13 that encodes an E3 ubiquitin ligase, but it is presently not known if this protein participates in the proteosomal degradation of C1GalT1. This signal will require further follow up. Importantly, our study demonstrates that there are several common genetic variants with relatively large effects on IgA1 *O*-glycosylation. These effects are conveyed by different genes, but converge on a single enzymatic step in the *O*-glycosylation pathway.

Our results contribute new insights into the genetic regulation of *O*-glycan synthesis, and demonstrate that a simple lectin-based assay can be used effectively to map genetic regulators of *O*-glycosylation of serum proteins. Given the high heritability of this trait, it is likely that additional loci contribute to variation in Gd-IgA1 levels. In particular, the inheritance pattern in IgAN kindreds suggested segregation of a major dominant gene, suggesting a potential role of additional rare alleles with large effects^11^. A search in larger population-based studies that includes both common and rare variants is likely to uncover additional genetic determinants of *O*-glycosylation defects and elucidate mechanisms leading to IgA nephropathy and related disorders.

## Materials and Methods

### Study Design and Power Analysis

The study was designed in two stages (**Supplementary Figure 1**). Stage 1 (the discovery phase) involved a genome-wide meta-analysis of two discovery cohorts: the Chinese cohort of 950 individuals (483 cases and 467 controls, all Han Chinese ancestry, genotyped with Illumina 660-quad chip), and the US cohort of 245 individuals (141 cases and 104 controls, all European ancestry, genotyped with the Illumina 550v3 chip). Genome-wide scan was performed in both cohorts and fixed-effects meta-analysis was applied to prioritize signals for follow-up studies. Stage 2 (the replication phase) involved genotyping of the top signals from stage 1 in five additional cohorts of European and Asian ancestry (1,438 individuals in total, Table 1). We carried out power calculations for this design and for a range of effect sizes under the following assumptions: standard normal trait distribution, additive risk model, no heterogeneity in association, marker allelic frequency of 0.25 (average MAF for the microarrays used), perfect LD between a marker and a causal allele, a follow-up significance threshold of P<5×10^−4^, and a joint significance level of P<5×10^−8^. These calculations demonstrate that we have adequate power to detect variants explaining >1.5% of overall trait variance (**Supplementary Table 3**). Our study was conducted according to the principles expressed in the Declaration of Helsinki; all subjects provided informed consent to participate in genetic studies, and the Institutional Review Board of Columbia University as well as local ethics review committees for each of the individual cohorts approved our study protocol.

### Phenotype Measurements and Quality Control

The level of serum total IgA was determined using standard ELISA^32^. The level of serum Gd-IgA1 was determined using custom HAA-based ELISA assay^11,12,32^. This method relies on the detection of HAA-lectin binding to desialylated galactose-deficient glycans (Tn antigens) of serum IgA1 immunocaptured on ELISA plates. Because in humans, IgA1, but not IgA2, has *O*-glycans, this assay effectively quantifies the absolute level of Gd-IgA1 in units/ml serum. We have optimized this assay for high-throughput use. Briefly, 96-well plates were coated with F(ab’)_2_ fragment of goat IgG anti-human IgA at 3 µg/ml. Plates were blocked with 1% BSA in PBS containing 0.05% Tween 20, and serial two-fold dilutions of samples and standards in blocking solution were incubated overnight at room temperature. To remove terminal sialic acid, the samples were treated with 100 µL (1 mU) per well of neuraminidase (Roche) in 10 mM sodium acetate buffer (pH=5) for 3 h at 37°C. Next, the samples were incubated with GalNAc-specific biotinylated HAA lectin (Sigma-Aldrich) for 3 h at 37°C. The bound lectin was detected with avidin-horseradish peroxidase conjugate, followed by the peroxidase substrate, o-phenylenediamine-H_2_O_2_ (Sigma); the reaction was stopped with. 1 M sulfuric acid. The concentration of Gd-IgA1 was calculated by interpolating the optical densities read at 490 nm on calibration curves constructed using a myeloma Gd-IgA1 standard. The intra-assay coefficients of variation (CVs) for calibration curves plotted by a 4-parameter model ranged from 2-10% for the extremes of the curves and 1-5% in the middle region. If higher values were noted, the samples were re-analyzed. The inter-assay CV was also consistently under 5% and our prior studies demonstrated excellent reproducibility of this assay^32^. In the final analysis, we applied a correction for potential plate effects using the same replicate samples across all plates. After corrections, serum Gd-IgA1 levels for each cohort were tested for normality by the Shapiro-Wilk test, assisted by visual inspection of histograms and qq-plots. Non-normal trait distributions were transformed using logarithmic transformation. The log-transformed traits were regressed against age and case-control status to derive standardized residuals. Summary statistics (mean, SD, skewness, and kurtosis) were derived for the distribution of standardized residuals, which were then used as a quantitative trait in GWAS analysis. Summary statistics, normality testing, transformations, plots, and regression analyses were performed with R 3.0 software package (CRAN).

### GWAS Discovery (Stage 1)

The genotyping, genotype quality control, and ancestry analyses of the discovery cohorts have been previously described^33,34^. Briefly, we implemented strict quality control filters for each cohort, eliminating samples with low call rates, duplicates, ancestry outliers, monomorphic and rare markers (MAF<1%), samples with cryptic relatedness, and samples with a detected sex mismatch. After all quality control steps, the Chinese Discovery Cohort was composed of 950 individuals typed with 508,112 SNPs, while the US Discovery Cohort was composed of 245 individuals typed with 531,778 SNPs. In total, 468,781 SNPs overlap between the cohorts. We used principal component–based ancestry matching algorithms to reduce any potential bias from population stratification (Spectral-GEM software)^35,36^, as described in prior studies of these cohorts^33,34^. Primary association testing for the Gd-IgA1 phenotype (expressed as standardized residuals) was performed within each cohort individually under an additive linear model in PLINK^37^. Significant principal components of ancestry were included as covariates in the association analysis of each individual cohort. Additionally, we performed regression analyses with and without adjustment for serum total IgA levels. Adjusted effect estimates and standard errors were derived for each SNP and the results were combined across the genome using an inverse variance-weighted method (METAL software)^38^. Genome-wide distributions of *P* values were examined visually using quantile-quantile plots for each individual cohort, as well as for the combined analysis. We estimated the genomic inflation factors^39^, which were negligible for each individual discovery cohort (lambda = 1.011 and 1.013 for the Chinese and US cohorts, respectively). The final meta-analysis QQ-plots showed no global departures from the expected distribution of *P* values and the overall genomic inflation factor was estimated at 1.010 (**Supplementary Figure 1**).

### Follow-up of Suggestive Signals (Stage 2)

We next prioritized the top 50 SNPs for replication among the top suggestive SNPs with P<5x10^−4^ from the GWAS analyses. First, we clustered the top signals into distinct loci based on their genomic coordinates and metrics of LD. Conditional regression analysis was carried out to detect independent association between signals within the same genomic regions. For genotyping in replication cohorts, we prioritized the independent SNPs with the lowest P-value in each independent locus. We additionally required that each SNP be successfully genotyped in both discovery cohorts. We also excluded loci supported by only a single SNP ('singleton signals' defined by the absence of supporting signals with P<0.01 within the same LD block). If genotyping of the top SNP failed, we selected a backup SNP on the basis of its strength of association, LD with the top SNP, quality of genotyping, and success in the design of working primers. Additionally, we added SNPs for which the signals became more significant (P<5×10^−4^) after adjustment for serum total IgA levels. In all, we successfully acquired and analyzed genotype data for 50 carefully selected SNPs in 1,438 independent replication samples across 5 cohorts. The ethnic composition of the replication cohorts, genotyping methods, and genotype call rates are summarized in Table 1. Association analyses were first carried out individually within each of the cohorts using the same methods as in the discovery study. The results were next combined using a fixed-effects model (**Supplementary Table 2**). For each SNP, we derived pooled effect estimates, their standard errors, and 95% confidence intervals. To declare genome-wide significance, we used the generally accepted threshold of P<5×10^−8^, initially proposed for Europeans genotyped with high-density platforms based on extrapolation to infinite marker density^40^.

### Chromosome X Analysis

We performed two types of association tests for X-linked markers. Our primary association test involved sex-stratified meta-analysis of chromosome X markers: each male and female sub-cohort was analyzed separately and the association statistics were combined across all sub-cohorts using fixed effects meta-analysis. This approach is not affected by the type of allele coding in males and allows for different effect size estimates between males and females^26^. In secondary analyses, we assumed complete X-inactivation in females and a similar effect size between males and females. In this test, females are considered to have 0, 1, or 2 copies of an allele as in an autosomal analysis while males are considered to have 0 or 2 copies of the same allele (*i.e.,* male hemizygotes are equivalent to female homozygotes). The main limitation of this approach relates the assumption of complete X inactivation. Because approximately 15-25% of X-linked genes escape inactivation in female-derived fibroblasts^41^ and chromosome X inactivation has not been studied in IgA1-secreting cells, this analysis was performed only on an exploratory basis, but the results were consistent with sex-stratified analyses.

### Tests of Alternative Inheritance Models and Epistasis

For the genome-wide significant loci, we explored two alternative genetic models (dominant and recessive) and compared these models using Bayesian Information Criterion (**Supplementary Table 8**). We also tested for all pairwise genetic interactions between the suggestive and significant loci using two different tests. First, we used a 1-degree-of-freedom likelihood ratio test (LRT) comparing two nested linear regression models: one with main effects only, and one with main effects and additive interaction terms. Second, we performed a more general 4-df genotypic interaction test. In this test, we compared a model with allelic effects, dominant effects, and their interaction terms with a reduced model without any of the interaction terms. All models were stratified by sex and cohort. The analyses were performed in R 3.0 software package (CRAN).

### Functional Annotation of Significant and Suggestive Loci

To interrogate putative functional SNPs that were not typed in our dataset, we systematically identified all variants that were in high LD (r^2^ > 0.5) with our top SNPs based on 1000 Genomes data. These variants were further annotated using ANNOVAR^42^, SeattleSeq^43^, SNPNexus^44^, FunciSNP^45^, HaploReg4^46^, and ChroMos^47^. Additionally, we identified all genes whose expression was correlated with the top SNPs in cis- or trans-using the following eQTL datasets: (1) meta-analysis of transcriptional profiles from peripheral blood cells of 5,311 Europeans^48^, (2) primary immune cells (B-cells and monocytes) from 288 healthy Europeans^49^, and (3) the latest release of GTEx data across multiple tissue types^19,50^. We utilized, automated MEDLINE text mining tools to assess network connectivity between genes residing in implicated GWAS loci, including GRAIL^51^, e-LiSe^52^, and FACTA+^53^. We also interrogated all known protein-protein interaction networks for connectivity between candidate genes using the Disease Association Protein-Protein Link Evaluator (DAPPLE)^54^ and Protein Interaction Network Analysis platform (PINA2)^55^. Network graphs were visualized in Cytoscape version 2.8.

### siRNA Knock-down Studies in IgA1 Secreting Cell Lines

IgA1-secreting cell lines from five patients with IgAN and five healthy controls were transfected using ON-TARGETplus SMARTpool siRNAs (Thermo Fisher Scientific, Lafayette, CO, USA) specific for human *C1GALT1, COSMC,* or both. The ON-TARGETplus Non-targeting Pool siRNAs was used as a control. We followed our previously published protocol for Amaxa nucleofector II (Lonza, Allendale, NJ, USA)^56^. Twenty-four hours after transfection, the knock-down efficiency was determined by qRT-PCR with previously described primers^8,56^. The knockdown was expressed as cDNA level of the individual gene normalized to GAPDH after respective siRNA treatment, divided by the respective value obtained after treatment by non-targeting siRNA. The effect of siRNA knock-down on the phenotype (the degree of galactose-deficiency of IgA1) was based on the reactivity of secreted IgA1 with a lectin from *Helix aspersa* specific for terminal GalNAc, as described^8,56^.

## Acknowledgements

We are grateful to all study participants for their contribution to this work. This study was supported by the following NIH grants from the National Institute for Diabetes and Digestive Kidney Diseases (NIDDK): K23DK090207 (K.K.), R03DK099564 (K.K.), R01DK105124 (K.K.), K01DK106341 (C.R.), R01DK078244 (J.N.), and R01DK082753 (A.G.G., J.N.), and by the Center for Glomerular Diseases at Columbia University.

